# Neuron-intrinsic NF-κB Signaling Mediates Reovirus Virulence

**DOI:** 10.1101/2020.08.05.238220

**Authors:** Andrea J. Pruijssers, Gwen Taylor, Pamela Brigleb, Pengcheng Shang, Kelly Urbanek, Judy J. Brown, Terence S. Dermody

## Abstract

Pathological effects of apoptosis associated with viral infections of the central nervous system are an important cause of morbidity and mortality. Reovirus is a neurotropic virus that causes apoptosis in neurons, leading to lethal encephalitis in newborn mice. Reovirus-induced encephalitis is diminished in mice with germline ablation of NF-κB subunit p50. It is not known whether the pro-apoptotic function of NF-κB is mediated by neuron-intrinsic processes, NF-κB-regulated cytokine production by inflammatory cells, or a combination of both. To determine the contribution of cell type-specific NF-κB signaling in reovirus-induced neuronal injury, we established mice that lack NF-κB p65 expression in neurons using the *Cre/loxP* recombination system. Following intracranial inoculation of reovirus, 50% of wild-type (WT) mice succumbed to infection, whereas more than 90% of mice lacking neural NF-κB p65 (Nsp65^−/−^) mice survived. While viral loads in brains of WT and Nsp65^−/−^ were comparable, histological analysis revealed that reovirus antigen-positive areas in the brain of WT mice displayed enhanced cleaved caspase-3 immunoreactivity, a marker of apoptosis, compared with Nsp65^−/−^ mice. These data suggest that neuron-intrinsic NF-κB-dependent factors are essential mediators of reovirus neurovirulence. RNA sequencing analysis of reovirus-infected cortices of WT and Nsp65^−/−^ mice suggests that NF-κB activation in neurons upregulates genes involved in innate immunity, inflammation, and cell death following reovirus infection. A better understanding of the contribution of cell type-specific NF-κB-dependent signaling to viral neuropathogenesis could inform development of new therapeutics that target and protect highly vulnerable cell populations

## INTRODUCTION

Many neurotropic viruses activate the NF-κB pathway during infection, which can contribute to cell survival and evasion of the immune response or elicit apoptosis and mediate viral spread (1). The role of NF-κB signaling in the central nervous system (CNS) is cell-type dependent. In neurons, NF-κB signaling maintains overall neuronal homeostasis, synapse growth and function, and neuroprotection under disease conditions. In glial cells, NF-κB signaling also contributes to maintaining neuronal cell health, although chronic NF-κB activation is neurotoxic (2). The factors that determine cell fate following viral induced-NF-κB activation in the CNS are not well understood.

Mammalian orthoreoviruses (reoviruses) are nonenveloped viruses with a segmented, double-stranded RNA genome that infect a variety of mammalian species but cause disease only in the very young. Reovirus is transmitted via the fecal-oral route and disseminates from the intestine to virtually all organs via hematogenous or neural routes. Infection of newborn mice with serotype 3 reovirus strains causes neuronal apoptosis, which leads to lethal encephalitis. Following cell entry, reovirus activates NF-κB using a mechanism that requires IKKα and IKKγ ((3)) and subsequent nuclear translocation of the canonical p50 and p65 NF-κB subunits ((3, 4)). NF-κB activation results in the expression of interferon-stimulated genes (ISGs), cytokines, chemokines, and other genes involved in antiviral responses and cell proliferation (5). Mouse embryo fibroblasts (MEFs) lacking either p50 or p65 display reduced levels of apoptosis following reovirus infection compared with wild-type (WT) MEFs (4), indicating that NF-κB subunits p50 and p65 mediate reovirus-induced apoptosis.

The role of p50 in reovirus virulence in newborn mice is organ-specific. In mice with germline ablation of NF-κB subunit p50 (p50^−/−^ mice), reovirus neurovirulence and neuronal apoptosis are diminished relative to WT mice. In contrast, p50^−/−^ mice succumb to severe myocarditis coupled with increased viral replication in the heart (6). These results suggested that NF-κB activation produces different outcomes depending on the cell type infected, a pro-apoptotic function in neurons versus an anti-apoptotic function in the heart. It is unclear whether the pro-apoptotic function of NF-κB is mediated by neuron-intrinsic processes, a result of NF-κB-regulated cytokine production by inflammatory cells, or a combination of both.

In this study, we established mice lacking NF-κB in neurons and investigated reovirus neuropathogenesis in these animals and WT mice. We discovered that mice with a neuron-specific deletion in the NF-κB p65-encoding gene (Nsp65^−/−^ mice) display increased survival following reovirus infection relative to WT mice and are likewise protected from brain injury. Brain regions in WT mice that harbor reovirus antigen also contain the cleaved (activated) form of the executioner apoptotic caspase, caspase-3. However, cleaved caspase-3 staining is markedly reduced in brains of Nsp65^−/−^ mice. To elucidate how NF-κB signaling in the CNS contributes to reovirus neuropathogenesis, we compared reovirus-induced transcriptional changes in brains of WT and Nsp65^−/−^ mice. Genes involved in innate immunity, inflammation, and cell death were identified as NF-κB pathways upregulated in the murine brain upon reovirus infection. These data identify potential therapeutic targets for development of antiviral agents to limit virus-induced neural injury.

## RESULTS

### Establishment of neuron-specific p65-deficient mice

To determine the contribution of neuron specific NF-κB signaling in reovirus-induced encephalitis *in vivo*, we crossed mice expressing Cre recombinase under control of the rat nestin promoter and enhancer to mice with loxP sites flanking exons 5 - 8 of p65/RelA (p65 ^f/f^) (7). Nestin is expressed in neuronal and glial precursor cells. Mice that carry the nestin Cre transgene and homozygous for the p65 floxed allele (neuron-specific p65-deficient [Nsp65^−/−^] mice) were present in the third generation (Fig. 1 A). Immunoblot analysis revealed that p65 expression was diminished in brains of Nsp65^−/−^ but not in WT p65 ^f/f^ mice (Fig. 1 B). Expression of p65 was not diminished in tissues that do not express nestin, demonstrating that p65 expression is conditionally ablated in brain tissue of Nsp65^−/−^ mice.

**Fig. 1.**
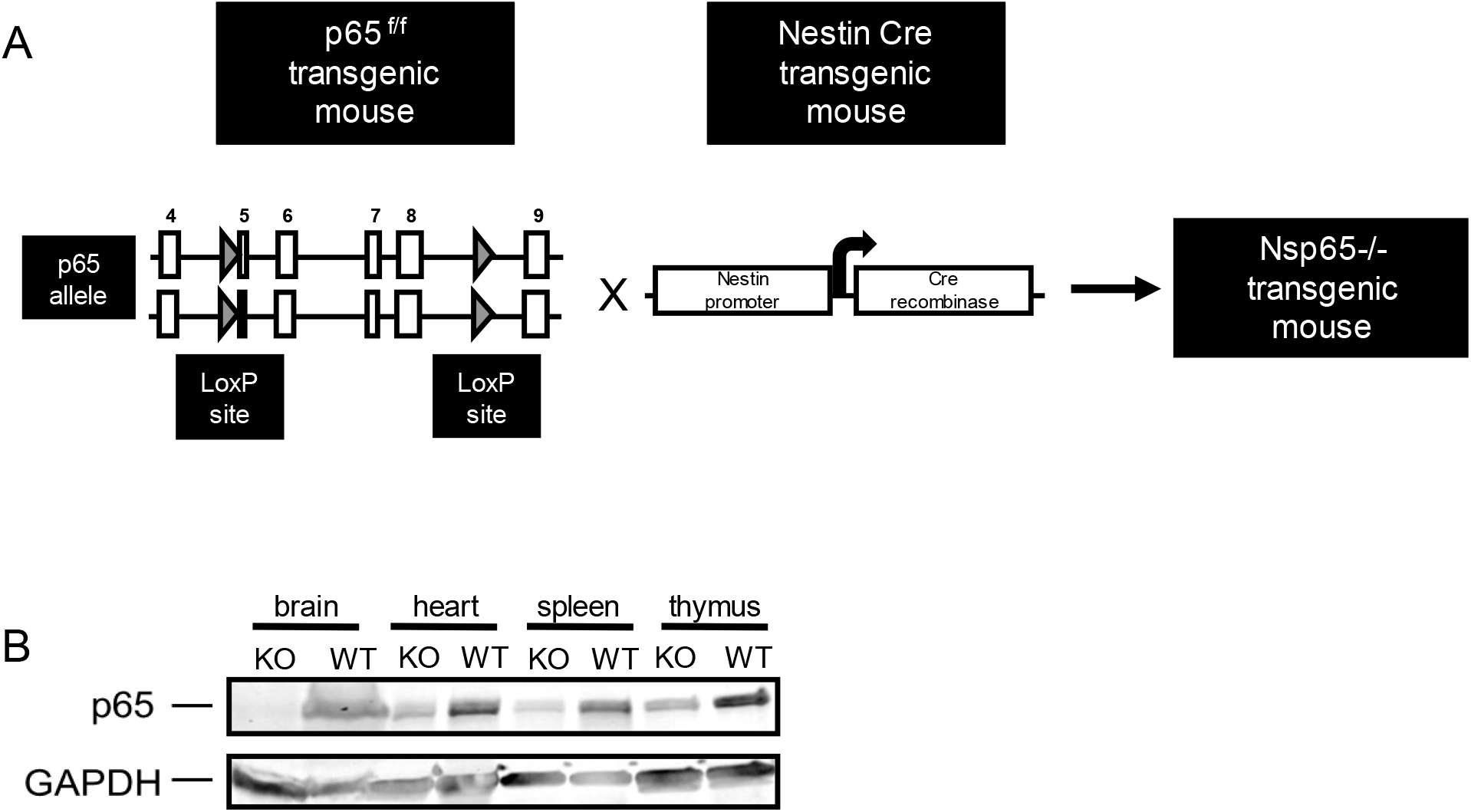
Neuron-specific p65-null mice lack p65 expression in the brain. (A) Breeding scheme for establishment of neuron-specific p65-null mice (Nsp65^−/−^). Mice with loxP sites (gray arrows) flanking exons 5 – 8 of the p65 allele (p65 ^f/f^) were interbred with mice expressing Cre recombinase under control of the rat nestin promoter and enhancer to establish mice heterozygous for the nestin Cre transgene and homozygous for the p65 floxed allele (Nsp65^−/−^). (B) Tissue homogenates from wild-type (WT) and Nsp65^−/−^ (KO) mice were resolved by SDS-PAGE and immunoblotted using antibodies specific for p65 and GAPDH.

### Apoptosis induced by reovirus infection is reduced in Nsp65^−/−^ cortical neurons

To determine whether neuron-intrinsic NF-κB-dependent factors contribute to reovirus-induced apoptosis, we infected cultures of WT, p50-deficient (p50^−/−^), and Nsp65^−/−^ cortical neurons with potently apoptotic reovirus strain T3SA+ and quantified the percentage of apoptotic neurons following acridine orange staining (Fig. 2 A, B). A significantly higher percentage of WT neurons were observed to undergo apoptosis following reovirus infection compared with p50^−/−^ and Nsp65^−/−^ neurons, suggesting that the pro-apoptotic function of NF-κB in the CNS is at least in part determined by neuron-intrinsic factors.

**Fig. 2.**
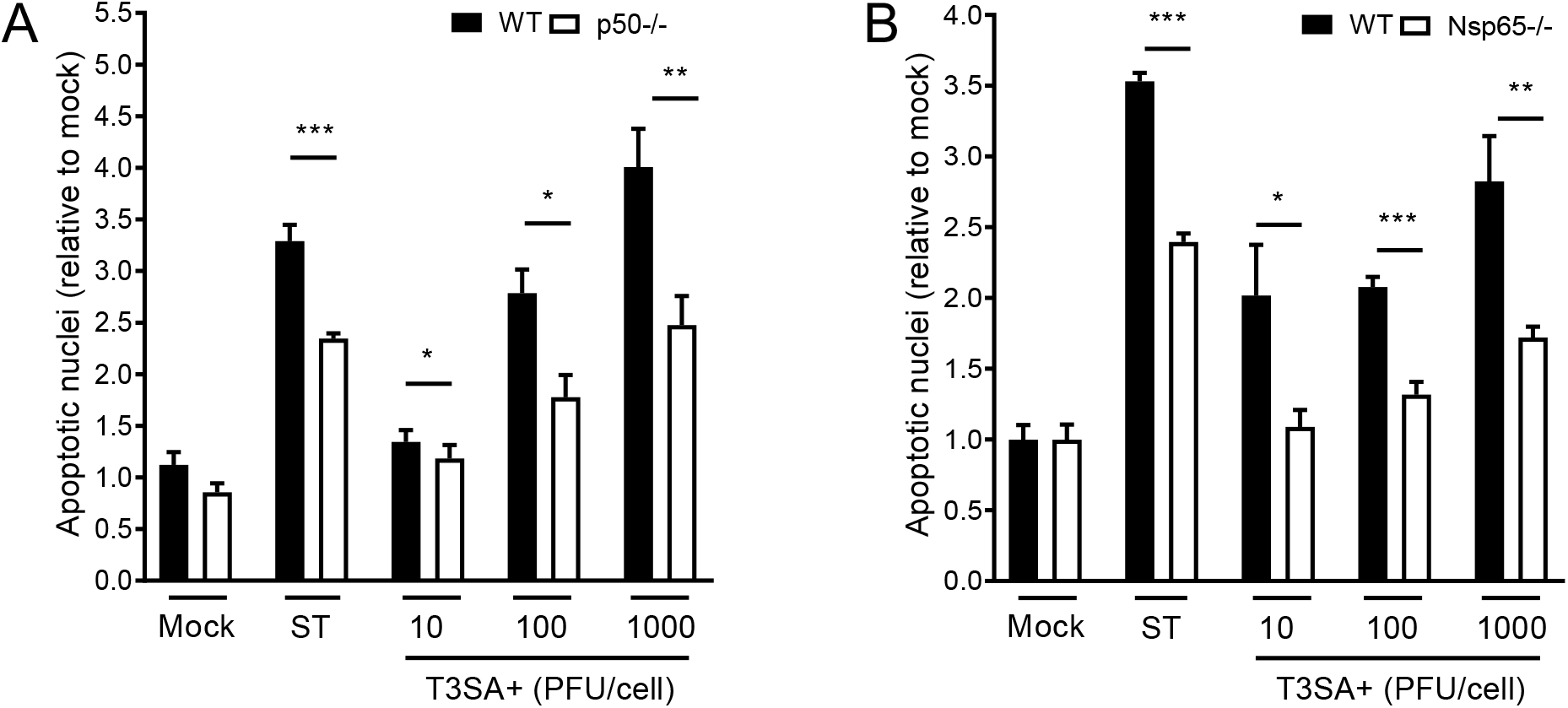
Reovirus-induced apoptosis is diminished in cultures of cortical neurons that lack either NF-κB p50 or p65. Wild-type (WT), p50^−/−^ (A), and Nsp65^−/−^ (B) cortical neurons were adsorbed with T3SA+ reovirus at the MOIs shown. At 24 h post-adsorption, the percentage of apoptotic nuclei were quantified following AO staining. Staurosporine (ST, 10 mM) was used as positive control. Each point is expressed as the mean percentage of apoptotic cells for triplicate wells. Error bars indicate SD. *, *P* < 0.05; **, *P* < 0.01; ***, *P* < 0.001 as determined by Student’s *t* test compared with results for Nsp65^−/−^ at the same MOI.

### Expression of NF-κB in neurons is required for reovirus virulence

To determine whether neuron-specific p65 contributes to reovirus neurovirulence, newborn WT and Nsp65^−/−^ mice were inoculated intracranially with T3SA+ and monitored for survival for 21 d (Fig. 3 A). While more than 90 percent of Nsp65^−/−^ mice survived T3SA+ infection, only 50 percent of WT mice survived, suggesting that p65 expression in neurons enhances reovirus neurovirulence.

**Fig. 3.**
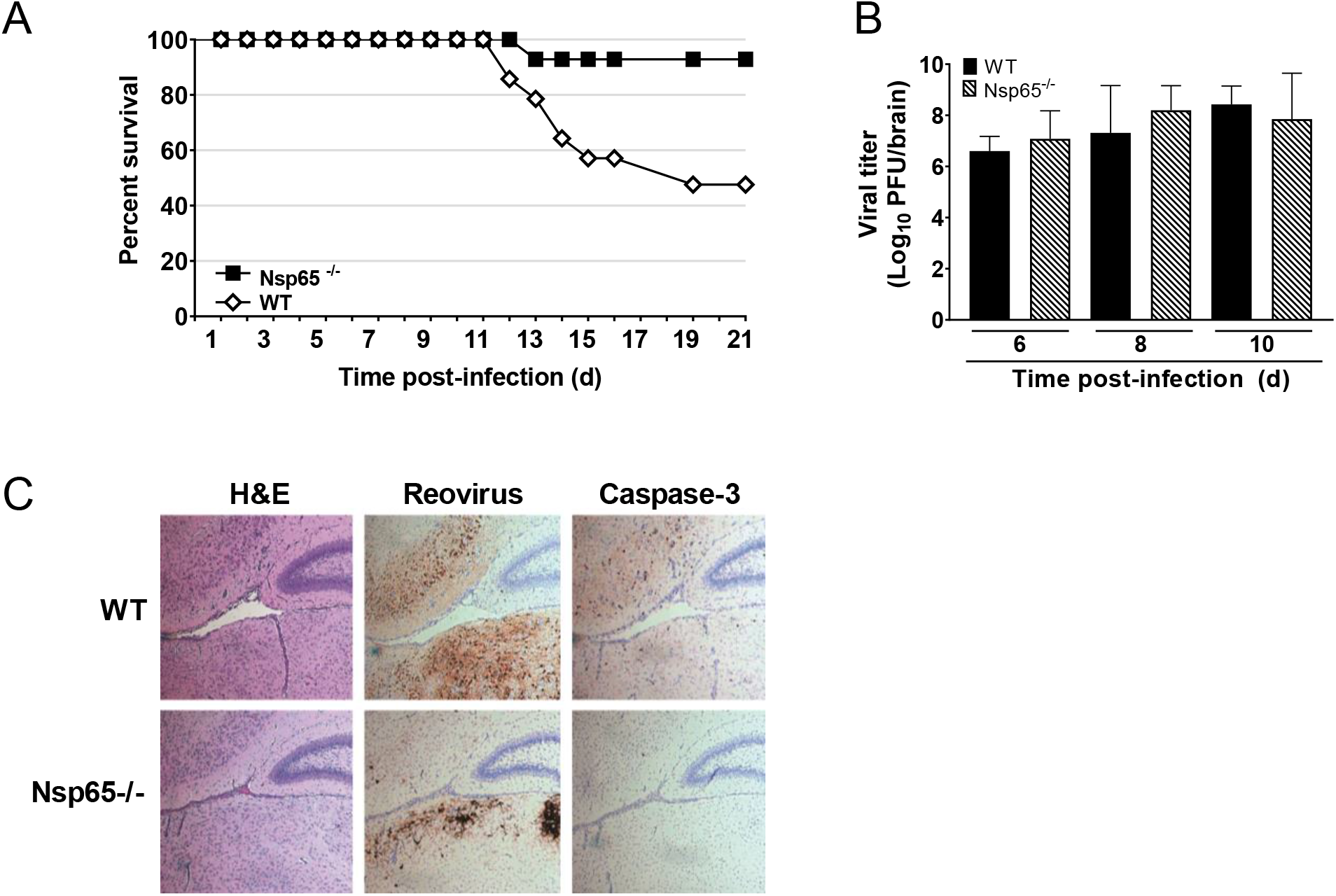
Neuron-specific NF-κB p65 is required for reovirus neurovirulence but not virus replication. (A) Survival following intracranial inoculation. Three-day-old wild-type (WT) and Nsp65^−/−^ mice (n = 14 WT and 16 Nsp65^−/−^) were inoculated intracranially with 10 PFU of reovirus T3SA+ and monitored for disease for 21 d. Mice were euthanized when moribund. Values that differ from WT by log rank test are indicated (*, *P* < 0.05). (B) Viral titers and (C) immunohistopathology in the brain following intracranial inoculation. An independent cohort of identically inoculated mice (n = 5 to 6 mice for each strain) were euthanized at 6, 8, and 10 d post-inoculation. (B) Viral titers of the homogenized left brain hemisphere were determined by plaque assay. Data are log transformed and displayed with a linear *x* axis scale. (C) Titer-matched right brain hemispheres at 6 d post-inoculation were fixed in formalin and embedded in paraffin. Coronal sections were either stained with H&E or immunostained with reovirus polyclonal antiserum or antibody specific for cleaved caspase-3. Representative sections are shown.

To determine whether neuron-specific NF-κB influences reovirus replication and pathology, newborn WT and Nsp65^−/−^ mice were inoculated intracranially with T3SA+. At 6, 8, and 10 d post-inoculation, mice were euthanized, and brains were resected. Viral loads detected in the left hemispheres of WT and Nsp65^−/−^ mice were comparable at all time points tested (Fig. 3 B). Serial coronal sections from embedded, titer-matched right hemispheres at 6 d post-inoculation were stained with hematoxylin and eosin (H&E) or immunostained with reovirus polyclonal antiserum or an antibody specific for the cleaved form of caspase-3 to monitor for cell death (Fig. 3 C). Areas in the brains of WT mice positive for reovirus antigen displayed enhanced cleaved caspase-3 immunoreactivity, whereas reovirus antigen-positive areas in the brains of Nsp65^−/−^ mice displayed low levels of cleaved caspase-3 immunoreactivity. Together, these data suggest that neuron-intrinsic NF-κB-dependent factors are not required for reovirus replication in the brain but are essential mediators of reovirus neurovirulence.

### NF-κB-dependent changes in mRNA expression in neurons following reovirus infection

To identify mediators of NF-κB-dependent, reovirus-induced apoptosis in neurons, we inoculated two-day-old WT and Nsp65^−/−^ mice intracranially with reovirus T3SA+ or PBS (mock). At 2 and 6 d post-inoculation, mice were euthanized, and cortices were micro-dissected, homogenized, and processed for viral titer via plaque assay and RNA purification. Transcript abundance in samples from infected WT and Nsp65^−/−^ mice, matched for viral titer, as well as those from PBS-inoculated controls was determined by mRNA sequencing. Three samples of RNA for each genotype per time point were subjected to paired-end RNA sequencing. We identified genes that were significantly up or down regulated in a reovirus- (897 genes), p65- (140 genes), and a reovirus- and p65- (814 genes) dependent manner and ranked the genes for abundance using DESeq2 (see methods) (Fig. 4A). The top 790 genes were uploaded to Ingenuity Software (Qiagen, Inc), which considers the adjusted *p* values of the analyses, and gene networks containing more than one candidate in the analysis were identified (Fig. 4B). Examples of gene networks that contained multiple candidates were related to inflammation, antimicrobial response, immune cell development, and cell death (Fig. 4B). Representative genes under direct control of p65 that were differentially induced in infected brain tissue were predominantly in groups involved in innate immunity, inflammation and cell death (Fig. 4C).

**Fig. 4.**
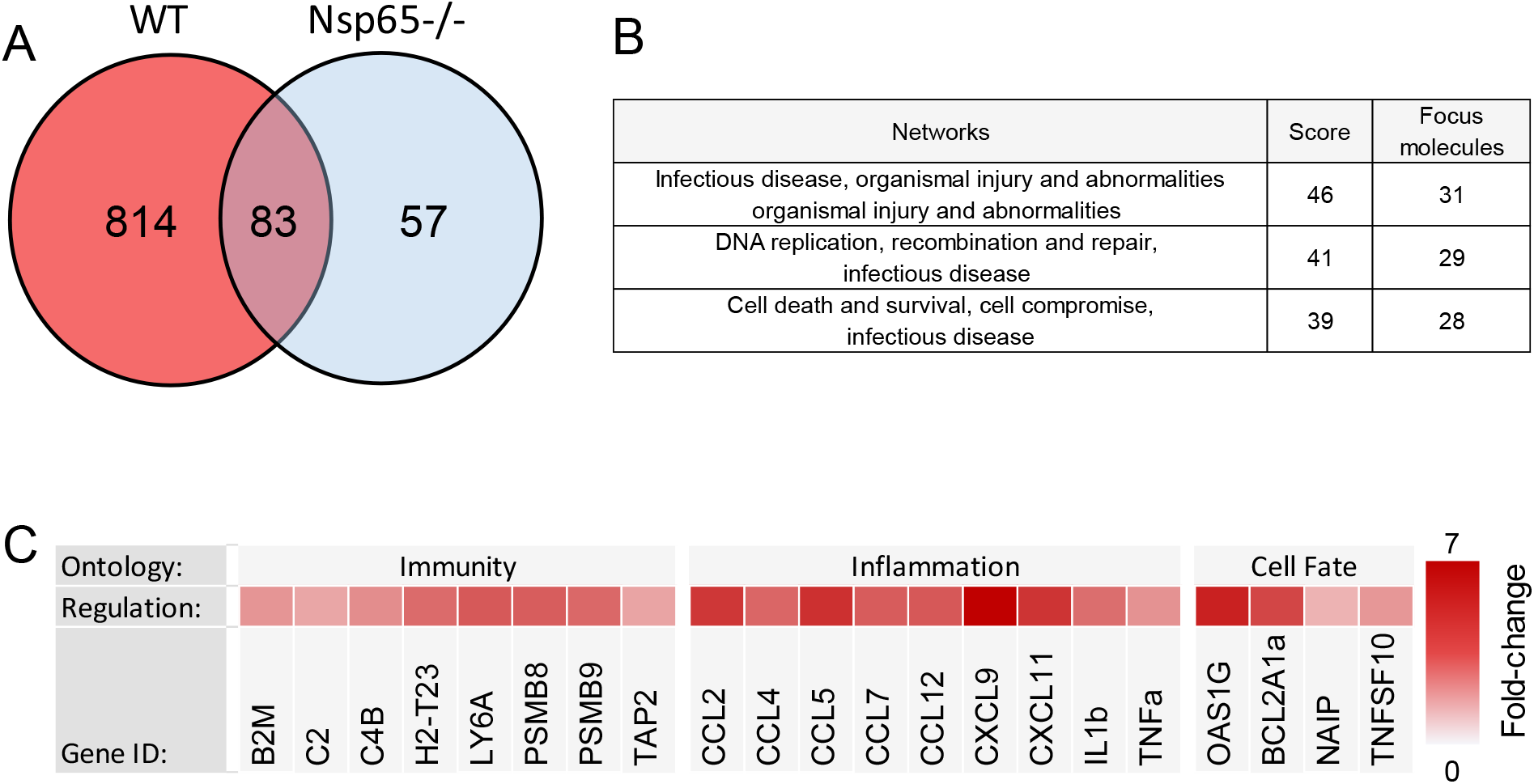
Expression of genes influenced by reovirus infection and NF-κB p65 in the brain. Wild-type (WT) and Nsp65^−/−^ newborn mice were inoculated intracranially with 10 PFU of reovirus T3SA+ or PBS. Cortices were removed at 6 d post-inoculation and processed for RNA purification and mRNA sequencing. RNA samples prepared from infected mice were matched for viral titer. (A) Venn diagram for differential expression analysis using DeSeq2 package in R between WT infected vs. mock (red) and Nsp65^−/−^ infected vs. mock (blue). (B) Networks with the most differentially expressed genes were identified using Ingenuity Pathway Analysis. (C) Heat map depicting expression levels.

### Reovirus infection in the brain induces a p65-dependent upregulation of proinflammatory cytokines

To validate at the protein level the pro-inflammatory cytokines that were upregulated at the transcript level in the presence of neuronal p65 (Fig. 4C), three-day-old WT and Nsp65^−/−^ mice were inoculated intracranially with T3SA+ or PBS. At 6 d post-inoculation, brains were harvested, homogenized, and processed for Luminex and plaque assay. Brain homogenates from WT and Nsp65^−/−^ mice, matched for viral titer, were analyzed for CCL2, CCL4, CCL5, CCL7, CCL12, and CXCL11 protein levels (Fig. 5). Levels of CCL2, CCL4, CCL5, CCL7, CCL12, and CXCL11 were upregulated in a reovirus- and p65-dependent manner, with statistically significant differences between WT and Nsp65^−/−^ mice observed for CCL4, CCL5, and CXCL11. These data validate our gene expression data and demonstrate that cytokine production during reovirus infection is dependent on NF-κB expression in neurons.

**Fig. 5.**
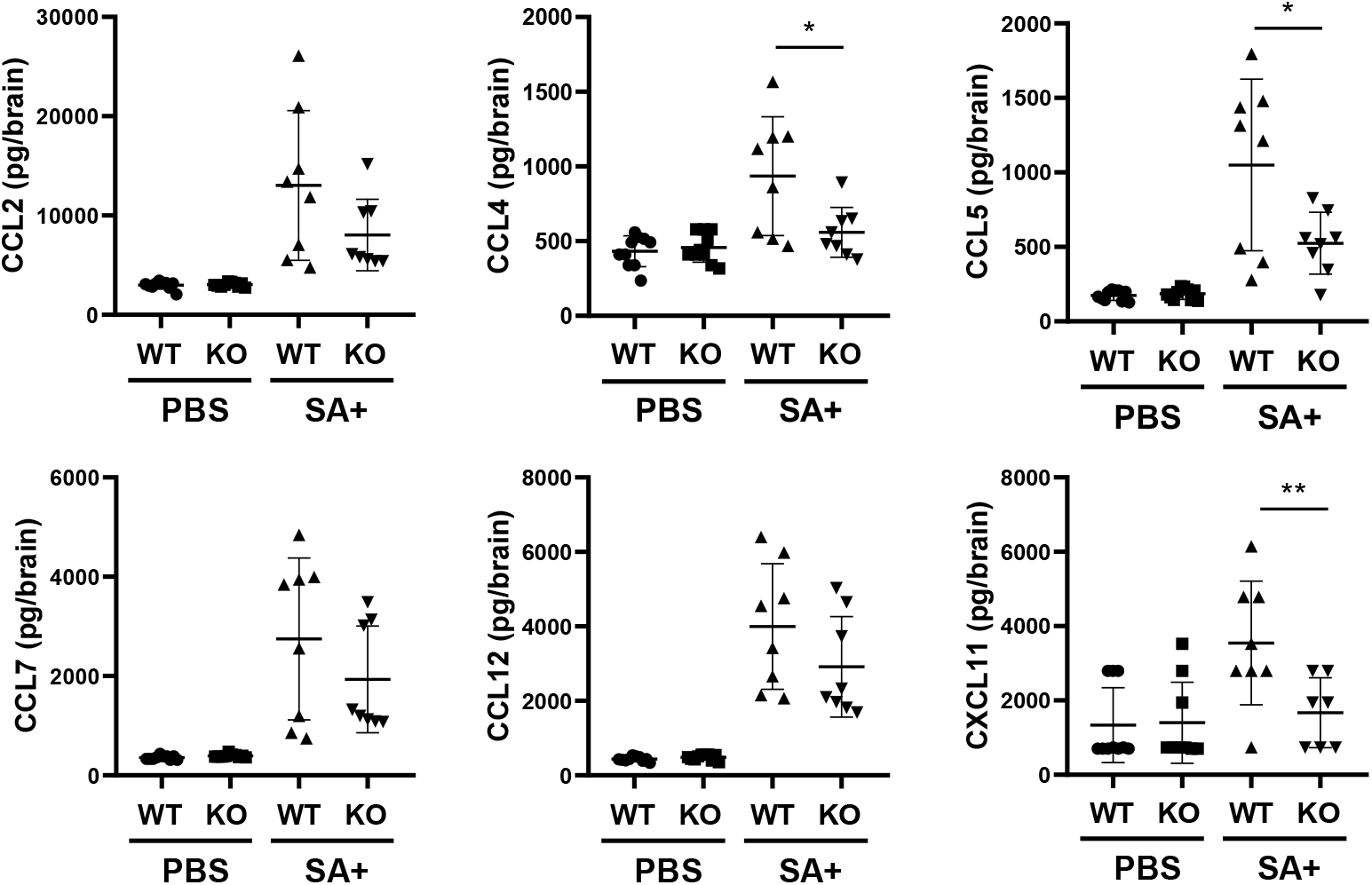
Reovirus-induced cytokine expression in the brain is dependent on NF-κB p65 expression in neurons. Wild-type (WT) and Nsp65^−/−^ (KO) newborn mice were inoculated intracranially with 10 PFU of reovirus T3SA+ or PBS. Cortices were removed at 6 d post-inoculation and processed for Luminex analysis and plaque assay. Samples were matched for viral titer. Data are presented as the mean values of 8 to 10 mice. *, *P* < 0.05; **, *P* < 0.005, as determined by unpaired Student’s *t* test compared with results for Nsp65^−/−^ mice.

### Proinflammatory cytokines are upregulated in reovirus-infected primary cortical neurons in an NF-κB dependent manner

While p65 expression is diminished in cells containing nestin in the brains of Nsp65^−/−^ mice, including neurons and microglia, p65 is expressed in cells that lack nestin, raising the possibility that surrounding cells in the brain contribute to the observed differences in cytokine production. To determine whether reovirus infection of neurons is sufficient to induce an increase in cytokine transcript levels in an NF-κB-dependent manner, we infected WT and Nsp65^−/−^ primary cortical neurons with T3SA+ infectious subvirion particles (ISVPs), a reovirus disassembly intermediate, that efficiently infects primary neuronal cultures (8). Transcript levels were detected by RT-PCR at 3, 6, 16, and 24 h post adsorption (Fig. 6). The level of S4 gene transcripts was used as a surrogate to quantify viral replication. At 6, 16, and 24 h post-adsorption, S4 gene transcript levels were significantly greater in infected Nsp65^−/−^ cortical neurons relative to WT cortical neurons, suggesting an NF-κB-dependent decrease in viral replication. However, we observed an NF-κB-dependent increase in transcript levels of *Ccl2*, *Ccl4*, *Ccl7*, and *Cxcl9*. Interestingly the increases in cytokine transcript levels can be divided into three groups based on kinetics of expression: *Ccl7* increased at 3 h post-adsorption and rapidly declined by 6 h post-adsorption (immediate early); *Ccl2* and *Ccl4* increased at 3 h post-adsorption but slowly declined over 13 h (early), and *Cxcl9* increased at 16 to 24 h post-adsorption (late). These data demonstrate that reovirus infection of primary cortical neurons upregulates cytokine gene expression in an NF-κB-dependent manner, suggesting that differences in cytokine levels observed in the brains of infected mice are at least in part attributable to NF-κB activation in neurons.

**Fig. 6.**
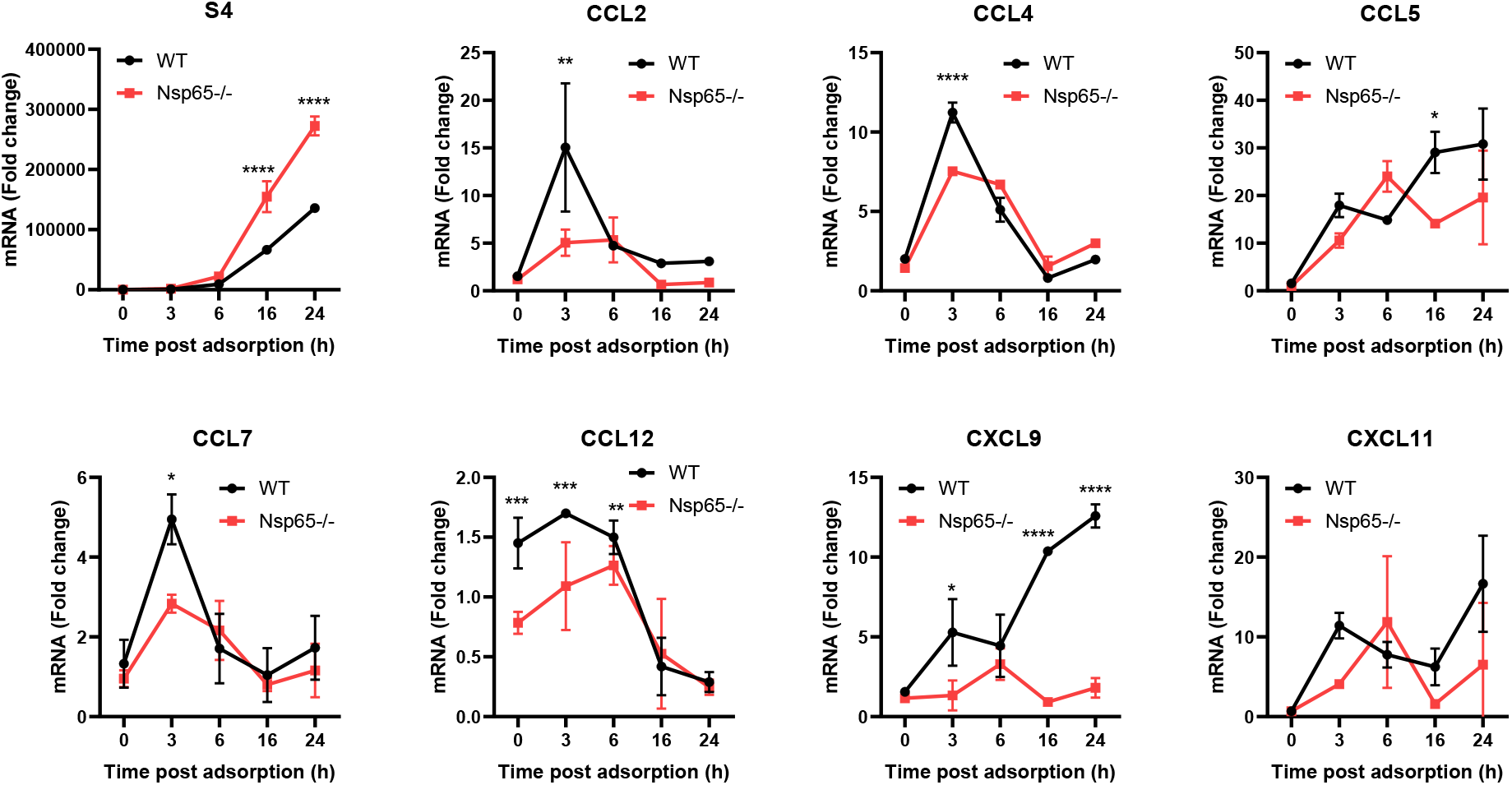
Reovirus-induced upregulation of cytokine transcripts in cultured cortical neurons is dependent on NF-κB p65 expression. Wild-type (WT) and Nsp65^−/−^ cortical neurons were adsorbed with 1000 PFU of reovirus T3SA+ ISVPs or PBS. Cells were harvested at times indicated, RNA was isolated, and transcript levels were determined by RT-qPCR. Data are presented as the mean values of two replicates. *, *P* < 0.05; **, *P* < 0.005; ****, *P* < 0.0001, as determined by two-way ANOVA with Sidak’s multiple comparisons.

## DISCUSSION

The NF-κB pathway is central to a variety of cellular processes including the regulation of cell cycle and cell fate, initiation of innate immune responses, and production of cytokines and chemokines. Several neurotropic viruses including enterovirus 71 (9, 10), herpes simplex virus-1 (HSV-1) (11), human immunodeficiency virus-1 (12), reovirus (4, 13), Sindbis virus (14), Venezuelan equine encephalitis virus (VEEV) (15), vesicular stomatitis virus (16), and West Nile virus (WNV) (17) activate NF-κB, which can lead to cell survival or apoptosis depending on the virus strain and cell type infected. Apoptosis induced by serotype 3 reovirus contributes to neuropathogenesis and is dependent on NF-κB activation (6). However, genes activated by NF-κB in response to reovirus infection in the CNS are unknown. In this study, we identified neuron-specific, reovirus-induced changes in the expression of genes under NF-κB control by establishing mice in which NF-κB p65 expression is ablated in neurons. We found that reovirus infection in the brain leads to upregulation of NF-κB-dependent host pathways involved in innate immunity, inflammation, and cell death.

Our previous studies using mice with germline deletion of NF-κB p50 indicate that NF-κB is required for reovirus to cause encephalitis but protects against reovirus myocarditis (6). To determine whether reovirus-induced neural injury is mediated by the activation of NF-κB in neurons, immune cells, or a combination of both, we established Nsp65^−/−^ mice in which NF-κB p65 is deleted in neurons (Fig 1). Nsp65^−/−^ mice displayed increased survival following reovirus infection relative to WT mice and were protected from severe brain injury (Fig. 3). These data suggest that neuron-intrinsic NF-κB-dependent factors are essential for reovirus neurovirulence.

An NF-κB-dependent gene network induced by reovirus infection in HeLa cells revealed temporal expression of distinct gene clusters, a majority of which are involved in innate immune responses and IFN signaling (5). However, genes activated by NF-κB can differ depending on the agonist or cell type (2, 18). We defined an NF-κB-dependent gene network induced by reovirus in the brain by RNA sequencing and observed that a majority of the genes identified are involved in innate immune responses and cell death (Fig. 4), similar to the findings made using HeLa cells (5).

In studies of a variety of neurotropic viruses, including dengue virus (19), Japanese encephalitis virus (JEV) (20), reovirus (21), VEEV (22), and WNV (20), the expression of genes encoding components of IFN signaling is most commonly altered (20). For example, JEV, reovirus, and WNV induce the expression of 86 common ISGs in the brain (20). These genes function in inflammation, immune signaling, virus recognition, and the antiviral response. Of these 86 ISGs common to neurotropic viral infections, we identified 72 that are upregulated during reovirus infection in WT mice. However, only 43 of these genes are expressed in response to NF-κB activation (Fig. 4). Interestingly, genes we identified under NF-κB control function in inflammation and immune signaling, while those that serve roles in viral recognition and the antiviral response are not NF-κB-dependent.

Neurotropic viral infections also induce elaboration of chemokines and recruitment of immune cells, which contribute to neuropathology (20, 23). We identified several chemokine genes that are expressed in response to reovirus infection and NF-κB activation, including *Ccl2*, *Ccl4*, *Ccl5*, *Ccl7*, *Ccl12*, *CxclL9*, and *Cxcl11* (Fig. 4). Furthermore, we found that chemokine levels also are upregulated in response to reovirus infection and NF-κB activation, with statistically significant increases observed for CCL4, CCL5, and CXCL11 (Fig. 5). Reovirus infection also upregulates CXCL10 both at the transcript and protein level, but this effect is independent of NF-κB expression (data not shown). Since transcript and protein levels were quantified at 6 d post-inoculation, it is possible that peak levels of CCL2, CCL7, and CCL12 are produced earlier in the infectious course and thus were not significantly increased in our analysis. Chemokines upregulated in the brain following reovirus infection also are upregulated by other neurotropic viruses. Following CNS infection, HSV-1 induces CCL2, CCL3, CCL5, CXCL9, and CXCL10, and JEV and WNV induce CCL2, CCL7, CXCL9, and CXCL10 (24). The similarities in chemokine responses elicited by neurotropic viruses suggest similar underlying pathological mechanisms and potentially common therapeutic targets.

Chemokines released during neurotropic viral infections can be produced by resident CNS cells or infiltrating immune cells (20, 25). Neurons are the primary source of chemokines following WNV infection, while astrocytes and microglia are the primary source following infection with HIV-1 (26) or HSV-1 (27). We hypothesized that neurons would be the primary source of chemokine production during reovirus infection, as these cells are heavily infected by reovirus in the brain. Using primary cortical neuron cultures prepared from WT and Nsp65^−/−^ mice, we found that transcript levels of *Ccl2*, *Ccl4*, *Ccl7*, *Ccl12*, and *Cxcl9* were increased in an NF-κB-dependent manner during reovirus infection, while those of *Ccl5* and *Cxcl11* were not. These data suggest that while reovirus-infected neurons contribute to the observed increase in chemokines in the brain, only a subset of these chemokines are dependent on NF-κB expression in neurons. Therefore, other cell types appear to modulate chemokine responses in the brain during reovirus infection. Since nestin is expressed in both neurons and microglia, NF-κB p65 is ablated in neurons and microglia in Nsp65^−/−^ mice. We think it possible that expression of *Ccl5* and *Cxcl11* in the CNS following reovirus infection is dependent on NF-κB activation in microglia rather than neurons.

Neurotropic viruses induce expression of highly orchestrated and distinct patterns of chemokines that recruit specific immune cell populations to sites of infection and contribute to disease. We observed that enhanced expression of the chemokine genes could be grouped into immediate-early (*Ccl7*), early (*Ccl2*, *Ccl4, Ccl12*), and late (*Cxcl9*) classes dependent on the kinetics of expression following reovirus infection of cortical neurons (Fig. 6). This time-dependent gene expression observed *in vitro* raises the possibility that expression of chemokines *in vivo* during reovirus infection of the CNS also is kinetically regulated. Understanding how this chemokine expression pattern is involved in immune cell infiltration and neuronal cell death is essential to understanding the role of NF-κB in reovirus neuropathogenesis.

Our studies enhance an understanding of the function of NF-κB in viral neuropathogenesis. We identified NF-κB-dependent pathways induced by reovirus infection in the CNS and discovered that expression of genes involved in immune signaling also are induced by other neurotropic viruses. Additionally, we identified a pattern of chemokine expression activated by reovirus in the brain that is in part recapitulated by reovirus-infected neurons. Collectively, these findings suggest that therapeutic interruption of the chemokine networks activated by reovirus may have broad utility against neurotropic viruses.

## MATERIALS AND METHODS

### Mice

Control p50^+/+^ (B6129PF1/J-A^W-J^/A^W^) and p50^−/−^ (B6129P-Nfkb^1tm1Bal^) mice (6, 28) and nestin Cre (B6.Cg-Tg(Nes-cre)1Kln/J) mice were obtained from Jackson Laboratory. RelA^F/F^ mice were provided by Dr. Albert Baldwin (29). Experiments were conducted using animal biosafety 2 (ABSL2) facilities and guidelines. All animal husbandry and experimental procedures were performed in accordance with U.S. Public Health Service policy and approved by the Institutional Animal Care and Use Committees at Vanderbilt University and the University of Pittsburgh.

### Establishment of neuron-specific p65-deficient mice

RelA^F/F^ (p65 ^f/f^) mice containing loxP sites flanking exons 5-8 of *RelA*, were interbred with mice expressing Cre recombinase under control of the rat nestin promoter and enhancer (Stock No: 003771: B6.Cg-Tg(Nes-cre)1Kln/J (30) to establish mice heterozygous for floxed *RelA* and the *nestin Cre* transgene (RelA^+/F^; nestin-Cre^+/-^). Mice carrying the *nestin Cre* transgene and homozygous for the *RelA* floxed allele (neuron-specific p65-deficient [Nsp65^−/−^] mice) were present in the third generation. For experimental breeding, Nsp65^−/−^ males were crossed with RelA^F/F^ females to establish litters with 50% of pups carrying the *nestin Cre* transgene (Nsp65^−/−^) and 50% without the *nestin Cre* transgene (WT). All experimental mice were genotyped with respect to *RelA* and *Cre* following necropsy.

### Cells and viruses

Spinner-adapted murine L929 (L) cells were maintained in Joklik’s modified Eagle’s minimal essential medium (US Biological, M3867) supplemented to contain 5% fetal bovine serum (FBS; VWR, 97068-085), 2 mM l-glutamine, 100 U/ml penicillin, 100 μg/ml streptomycin, and 25 ng/ml amphotericin B. Primary neuronal cultures were derived from cortices of p50^+/+^ (B6129PF1/J-A^W-J^/A^W^), p50^−/−^ (B6129P-Nfkb^1tm1Bal^), Nsp65^−/−^ (RelA^F/F^; nestin-Cre^+/-^), or WT (RelA^F/F^; nestin-Cre^−/−^) day 15 embryos as described (31). Neurons were plated at a density of 10^5^ cells/well in 24-well plates (Greiner) pre-treated with 10 mg/ml poly-D-lysine hydrobromide (Sigma-Aldrich, P0899) and 1.64 mg/ml laminin (Corning, 354232) diluted in Neurobasal medium (Gibco, 21103049). Cultures were incubated for 24 h in Neurobasal medium supplemented to contain 10% FBS, 0.6 mM GlutaMAX, 50 U/ml penicillin, and 50 μg/ml streptomycin. Cultures thereafter were maintained in Neurobasal medium supplemented to contain 1 × B-27 (Gibco, 35050079), 0.6 mM GlutaMAX, 50 U/ml penicillin, and 50 μg/ml streptomycin. One-half of the medium volume was replaced with fresh medium every 3 to 4 d. Neurons were cultivated for 5 to 7 d prior to use.

Purified reovirus strain T3SA+ was prepared from second- or third-passage L-cell lysate stocks (32). Viral particles were Vertrel XF-extracted from infected cell lysates, layered onto 1.2- to 1.4-g/cm^3^ CsCl gradients, and centrifuged at 62,000 × *g* for 16 h. Bands corresponding to virions (1.36 g/cm^3^) were collected and dialyzed in virion-storage buffer (150 mM NaCl, 15 mM MgCl_2_, 10 mM Tris-HCl [pH 7.4]) (33). The concentration of reovirus virions in purified preparations was determined from an equivalence of one OD unit at 260 nm equals 2.1 × 10^12^ virions (33). Viral titer was determined by plaque assay using L cells (34). ISVPs were prepared by treating 2 × 10^11^ virion particles with 20 μg of α-chymotrypsin (Sigma-Aldrich) in a 100-μl volume of virion storage buffer at 35°C for 60 min (35). Reactions were terminated by the addition of 2 mM phenylmethylsulphonyl fluoride (Sigma-Aldrich). ISVP titer was determined by plaque assay using L cells.

### SDS-PAGE and immunoblotting

Samples for SDS-PAGE were diluted in 5 × Laemmli sample buffer (BioRad) containing 10% β-mercaptoethanol, incubated at 95°C for 5 min, and resolved in 10% Mini-Protean TGX gels (Bio-Rad). Proteins were transferred to nitrocellulose membranes (Bio-Rad) at 100 V at 4°C for 1 h and immunoblotted using rabbit anti-p65 (Abcam, ab16502), cleaved caspase-3, (Cell Signaling Technology, 9664), and mouse anti-GAPDH (Sigma-Aldrich, G8795). Secondary antibodies IRDye® 680RD goat anti-rabbit IgG and IRDye® 800CW goat anti-mouse IgG (Li-Cor) were used for detection. Membranes were scanned using an Odyssey CLx imaging system (Li-Cor), and pixel intensities of the bands were quantified using Image Studio Software (Li-Cor).

### Quantification of apoptosis by acridine orange (AO) staining

Neurons were adsorbed with a multiplicity of infection (MOI) of 1000 reovirus ISVPs per cell diluted in PBS at 37°C for 1 h. The inoculum was removed, fresh medium was added, and neurons were incubated at 37°C for 24 or 48 h. The percentage of apoptotic cells was determined using AO staining as described (36). Images were collected from > 600 cells in two to three fields of view per well by epi-illumination fluorescence microscopy using a fluorescein filter set (Zeiss Photomicroscope III).

### Infection of mice

Two-to-three-day-old WT or Nsp65^−/−^ mice weighing 1.6 to 2.3 g were inoculated intracranially in the right cerebral hemisphere with 5 μl of 2 plaque-forming units (PFU)/μl of purified T3SA+ diluted in PBS using a 30-gauge needle and syringe (Hamilton). Titers of virus in the inoculum were determined to confirm the number of infectious particles in the final dilution. The results of these assays are reported as the inoculating viral dose. For analysis of virulence, mice were monitored for symptoms of disease for 21 d post-inoculation and euthanized when moribund, as defined by severe lethargy or seizures, paralysis, or loss of 25% of body weight. Death was not used as an endpoint in our experiments.

Viral replication and immunohistopathology were analyzed at 6 d post-inoculation. Mice were euthanized, and brains were removed and hemisected along the longitudinal fissure. The left hemisphere was collected in 1 ml PBS, frozen and thawed once, and homogenized using a TissueLyser (Qiagen, Inc.). Viral titers in brain homogenates were determined by plaque assay using L929 cells (37). The right hemisphere was submerged in 10% buffered formalin at room temperature for 24 to 120 h, embedded in paraffin, and cut into consecutive 6-μm sections. Consecutive sections were processed as described (37) for evaluation of histopathologic changes by H&E staining or for immunohistochemical detection of reovirus antigen (1:45,000 dilution) or the cleaved (active) form of caspase-3 (Cell Signaling, 9664; 1:300 dilution). Images were captured at a magnification of ×20 and a resolution of 0.5 μm/pixel using a high-throughput Leica SCN400 slide scanner automated digital imaging system.

### RNA sequencing

Two-to-three-day-old WT or Nsp65^−/−^ mice were inoculated intracranially with T3SA+ and euthanized at 2 and 6 d post-inoculation. Cortices were microdissected, homogenized, and processed for plaque assay and RNA purification. Tails were harvested for genotyping. Total RNA was extracted from brain homogenates using TRIzol, digested with DNase, and further purified using the High Pure RNA Isolation kit (Roche). RNA quantity was assessed using the Quant-iT RiboGreen RNA Assay kit (Invitrogen), and RNA quality was assessed using an Agilent bioanalyzer. A total of 1 mg of RNA was used for library preparation. Transcript abundance in samples from infected WT and Nsp65^−/−^ mice matched for viral titer as well as those from PBS-inoculated controls were determined by mRNA sequencing using a HiSeq 2000 instrument (Illumina). Three samples of RNA for each genotype per time point were subjected to paired-end RNA sequencing. Statistical analysis identified genes that were significantly increased or decreased in a reovirus-, p65-, and a reovirus- and p65-dependent manner.

### Data analysis

Quality-controlled FASTQ files were aligned to the Ensemble *Mus musculus* genome (GRCm38) using STAR aligner (version 2.5.1). HTSeq-count was used to quantify counts of reads uniquely mapped to annotated genes using the GRCm38 annotation gtf file (38). Differential gene expression by age, brain regions, p65 expression, and infection was analyzed using DESeq2 (39) using a model based on the negative binomial distribution. The resulting *P* values were adjusted using the Benjamini and Hochberg approach for controlling the false discovery rate, and differentially expressed genes were identified at the 5% threshold. Gene-set enrichment analysis was used to assess the statistical enrichment of gene ontologies and pathways (40).

### Luminex assays

Brain homogenates prepared from mice 6 d post-inoculation were prepared as described for viral titer. Lysates were analyzed by Luminex profiling (Luminex Corporation) using the Bio-Plex Pro Mouse Chemokine 31-plex panel (Bio-Rad; 12009159) at a 1:2 dilution following the manufacturer’s instructions. Cytokine abundance was determined using the laboratory multianalyte profiling system (LabMAP, Luminex).

### Infection of primary cortical neurons

Neurons were adsorbed with T3SA+ ISVPs at an MOI of 1000 PFU/cell at 37°C for 1 h, washed with PBS, and incubated at 37°C for various intervals. Cells were washed with PBS and lysed in 1 mL TRIzol for RNA isolation.

### qPCR and complementary DNA synthesis

Total RNA was extracted from infected cortical neurons at various intervals using TRIZOL and an RNeasy Mini kit (Qiagen) according to manufacturer’s instructions, followed by DNase digestion. RNA was reverse-transcribed with the SuperScript IV First-Strand Synthesis system (ThermoFisher) using 0.5 to 1 μg RNA. RT-qPCR was conducted using SsoAdvanced Universal Probes Supermix (BioRad) with an Applied Biosystems ViiA7. Gene expression was determined by the ΔΔC_q_ method in which cDNA abundance was normalized to mouse HPRT. Primers from Thermo Fisher include: CCL2 (Mm00441242_m1), CCL4 (Mm00443111_m1), CCL5 (Mm01302427_m1), CCL7 (Mm00443113_m1), CCL12 (Mm01617100_m1), CXCL9 (Mm00434946_m1), CXCL11 (Mm00444662_m1), TNFα (Mm00443258_m1), and HPRT (Mm00446968_m1).

### Statistical analysis

Experiments were performed in triplicate and repeated at least twice. Representative results of single experiments are shown. Mean values were compared using Student’s *t* test, Mann-Whitney test, Fisher’s Combined Probability test, 2-way ANOVA with Sidak’s multiple comparisons, or Log-rank test. Error bars denote the range of data or standard deviation (S. D.). *P* values of < 0.05 were considered to be statistically significant.

## Acknowledgments

We thank members of the Dermody laboratory for insightful discussions regarding this work. We thank Dr. Albert Baldwin at the University of North Carolina Chapel Hill for sharing with us the strain of mice containing floxed p65/RelA developed in his laboratory. We acknowledge the staff of the Vanderbilt Translational Pathology Shared Resource and the Research Pathology/Health Sciences Tissue Bank at UPMC Hillman Cancer Center.

This work was supported by the U.S. Public Health Service awards T32 AI60525 (P.B.), T32 HL007751 and F31 DK108562 (J.J.B.), and R01 AI038296 (T.S.D.) and the Elizabeth B. Lamb Center for Pediatric Research. The funders had no role in study design, data collection and analysis, decision to publish, or preparation of the manuscript.

## REFERENCES

1. Hiscott J, Kwon H, Genin P. 2001. Hostile takeovers: viral appropriation of the NF-kappaB pathway. J Clin Invest 107:143–51.

2. Dresselhaus EC, Meffert MK. 2019. Cellular Specificity of NF-κB Function in the Nervous System. Front Immunol 10:1043.

3. Hansberger MW, Campbell JA, Danthi P, Arrate P, Pennington KN, Marcu KB, Ballard DW, Dermody TS. 2007. IkB kinase subunits a and g are required for activation of NF-kB and induction of apoptosis by mammalian reovirus. Journal of Virology 81:1360–1371.

4. Connolly JL, Rodgers SE, Clarke P, Ballard DW, Kerr LD, Tyler KL, Dermody TS. 2000. Reovirus-induced apoptosis requires activation of transcription factor NF-kB. Journal of Virology 74:2981–2989.

5. O’Donnell SM, Holm GH, Pierce JM, Tian B, Watson MJ, Chari RS, Ballard DW, Brasier AR, Dermody TS. 2006. Identification of an NF-kB-dependent gene network in cells infected by mammalian reovirus. Journal of Virology 80:1077–1086.

6. O’Donnell SM, Hansberger MW, Connolly JL, Chappell JD, Watson MJ, Pierce JM, Wetzel JD, Han W, Barton ES, Forrest JC, Valyi-Nagy T, Yull FE, Blackwell TS, Rottman JN, Sherry B, Dermody TS. 2005. Organ-specific roles for transcription factor NF-kappaB in reovirus-induced apoptosis and disease. J Clin Invest 115:2341–50.

7. Steinbrecher KA, Harmel-Laws E, Sitcheran R, Baldwin AS. 2008. Loss of epithelial RelA results in deregulated intestinal proliferative/apoptotic homeostasis and susceptibility to inflammation. J Immunol 180:2588–2599.

8. Pruijssers AJ, Hengel H, Abel TW, Dermody TS. 2013. Apoptosis induction influences reovirus replication and virulence in newborn mice. J Virol 87:12980–9.

9. Tung WH, Lee IT, Hsieh HL, Yang CM. 2010. EV71 induces COX-2 expression via c-Src/PDGFR/PI3K/Akt/p42/p44 MAPK/AP-1 and NF-kappaB in rat brain astrocytes. J Cell Physiol 224:376–86.

10. Hsiao HB, Chou AH, Lin SI, Chen IH, Lien SP, Liu CC, Chong P, Liu SJ. 2014. Toll-like receptor 9-mediated protection of enterovirus 71 infection in mice is due to the release of danger-associated molecular patterns. Journal of Virology 88:11658–70.

11. Gregory D, Hargett D, Holmes D, Money E, Bachenheimer SL. 2004. Efficient replication by herpes simplex virus type 1 involves activation of the IkappaB kinase-IkappaB-p65 pathway. Journal of Virology 78:13582–90.

12. Liu R, Tan J, Lin Y, Jia R, Yang W, Liang C, Geng Y, Qiao W. 2013. HIV-1 Vpr activates both canonical and noncanonical NF-κB pathway by enhancing the phosphorylation of IKKα/β. Virology 439:47–56.

13. O’Donnell SM, Hansberger MW, Connolly JL, Chappell JD, Watson MJ, Pierce JM, Wetzel JD, Han W, Barton ES, Forrest JC, Valyi-Nagy T, Yull FE, Blackwell TS, Rottman JN, Sherry B, Dermody TS. 2005. Organ-specific roles for transcription factor NF-kB in reovirus-induced apoptosis and disease. Journal of Clinical Investigation 115:2341–2350.

14. Lin KI, Lee SH, Narayanan R, Baraban JM, Hardwick JM, Ratan RR. 1995. Thiol agents and Bcl-2 identify an alphavirus-induced apoptotic pathway that requires activation of the transcription factor NF-kappa B. Journal of Cell Biology 131:1149–61.

15. Amaya M, Voss K, Sampey G, Senina S, de la Fuente C, Mueller C, Calvert V, Kehn-Hall K, Carpenter C, Kashanchi F, Bailey C, Mogelsvang S, Petricoin E, Narayanan A. 2014. The role of IKKβ in Venezuelan equine encephalitis virus infection. PLoS ONE 9:e86745.

16. Boulares AH, Ferran MC, Lucas-Lenard J. 1996. NF-kappaB activation Is delayed in mouse L929 cells infected with interferon suppressing, but not inducing, vesicular stomatitis virus strains. Virology 218:71–80.

17. Kesson AM, King NJ. 2001. Transcriptional regulation of major histocompatibility complex class I by flavivirus West Nile is dependent on NF-kappaB activation. J Infect Dis 184:947–54.

18. Hiscott J, Kwon H, Génin P. 2001. Hostile takeovers: viral appropriation of the NF-kappaB pathway. J Clin Invest 107:143–51.

19. Bordignon J, Probst CM, Mosimann AL, Pavoni DP, Stella V, Buck GA, Satproedprai N, Fawcett P, Zanata SM, de Noronha L, Krieger MA, Duarte Dos Santos CN. 2008. Expression profile of interferon stimulated genes in central nervous system of mice infected with dengue virus Type-1. Virology 377:319–29.

20. Clarke P, Leser JS, Bowen RA, Tyler KL. 2014. Virus-induced transcriptional changes in the brain include the differential expression of genes associated with interferon, apoptosis, interleukin 17 receptor A, and glutamate signaling as well as flavivirus-specific upregulation of tRNA synthetases. mBio 5:e00902–14.

21. Tyler KL, Leser JS, Phang TL, Clarke P. 2010. Gene expression in the brain during reovirus encephalitis. J Neurovirol 16:56–71.

22. Sharma A, Bhattacharya B, Puri RK, Maheshwari RK. 2008. Venezuelan equine encephalitis virus infection causes modulation of inflammatory and immune response genes in mouse brain. BMC Genomics 9:289.

23. Chandwani MN, Creisher PS, O’Donnell LA. 2019. Understanding the Role of Antiviral Cytokines and Chemokines on Neural Stem/Progenitor Cell Activity and Survival. Viral Immunol 32:15–24.

24. Klein DDaR. 2011. Pathogenesis of Encephalitis: Chemokines and viral infections of the CNS.

25. Hosking MP, Lane TE. 2010. The role of chemokines during viral infection of the CNS. PLoS Pathog 6:e1000937.

26. van Marle G, Henry S, Todoruk T, Sullivan A, Silva C, Rourke SB, Holden J, McArthur JC, Gill MJ, Power C. 2004. Human immunodeficiency virus type 1 Nef protein mediates neural cell death: a neurotoxic role for IP-10. Virology 329:302–18.

27. Aravalli RN, Hu S, Rowen TN, Palmquist JM, Lokensgard JR. 2005. Cutting edge: TLR2-mediated proinflammatory cytokine and chemokine production by microglial cells in response to herpes simplex virus. J Immunol 175:4189–93.

28. Sha WC, Liou HC, Tuomanen EI, Baltimore D. 1995. Targeted disruption of the p50 subunit of NF-kappa B leads to multifocal defects in immune responses. Cell 80:321–30.

29. Steinbrecher KA, Harmel-Laws E, Sitcheran R, Baldwin AS. 2008. Loss of epithelial RelA results in deregulated intestinal proliferative/apoptotic homeostasis and susceptibility to inflammation. J Immunol 180:2588–99.

30. Luo L, Ambrozkiewicz MC, Benseler F, Chen C, Dumontier E, Falkner S, Furlanis E, Gomez AM, Hoshina N, Huang WH, Hutchison MA, Itoh-Maruoka Y, Lavery LA, Li W, Maruo T, Motohashi J, Pai EL, Pelkey KA, Pereira A, Philips T, Sinclair JL, Stogsdill JA, Traunmüller L, Wang J, Wortel J, You W, Abumaria N, Beier KT, Brose N, Burgess HA, Cepko CL, Cloutier JF, Eroglu C, Goebbels S, Kaeser PS, Kay JN, Lu W, Luo L, Mandai K, McBain CJ, Nave KA, Prado MAM, Prado VF, Rothstein J, Rubenstein JLR, Saher G, Sakimura K, Sanes JR, Scheiffele P, Takai Y, et al. 2020. Optimizing Nervous System-Specific Gene Targeting with Cre Driver Lines: Prevalence of Germline Recombination and Influencing Factors. Neuron 106:37–65.e5.

31. Antar AAR, Konopka JL, Campbell JA, Henry RA, Perdigoto AL, Carter BD, Pozzi A, Abel TW, Dermody TS. 2009. Junctional adhesion molecule-A is required for hematogenous dissemination of reovirus. Cell Host Microbe 5:59–71.

32. Furlong DB, Nibert ML, Fields BN. 1988. Sigma 1 protein of mammalian reoviruses extends from the surfaces of viral particles. Journal of Virology 62:246–56.

33. Smith RE, Zweerink HJ, Joklik WK. 1969. Polypeptide components of virions, top component and cores of reovirus type 3. Virology 39:791–810.

34. Virgin HW, IV, Bassel-Duby R, Fields BN, Tyler KL. 1988. Antibody protects against lethal infection with the neurally spreading reovirus type 3 (Dearing). Journal of Virology 62:4594–4604.

35. Baer GS, Dermody TS. 1997. Mutations in reovirus outer-capsid protein s3 selected during persistent infections of L cells confer resistance to protease inhibitor E64. Journal of Virology 71:4921–4928.

36. Tyler KL, Squier MK, Rodgers SE, Schneider SE, Oberhaus SM, Grdina TA, Cohen JJ, Dermody TS. 1995. Differences in the capacity of reovirus strains to induce apoptosis are determined by the viral attachment protein s1. Journal of Virology 69:6972–6979.

37. Sutherland DM, Aravamudhan P, Dietrich MH, Stehle T, Dermody TS. 2018. Reovirus neurotropism and virulence are dictated by sequences in the head domain of the viral attachment protein. J Virol 92:e00974–18.

38. Anders S, Pyl PT, Huber W. 2015. HTSeq--a Python framework to work with high-throughput sequencing data. Bioinformatics 31:166–9.

39. Love MI, Huber W, Anders S. 2014. Moderated estimation of fold change and dispersion for RNA-seq data with DESeq2. Genome Biol 15:550.

40. Subramanian A, Tamayo P, Mootha VK, Mukherjee S, Ebert BL, Gillette MA, Paulovich A, Pomeroy SL, Golub TR, Lander ES, Mesirov JP. 2005. Gene set enrichment analysis: a knowledge-based approach for interpreting genome-wide expression profiles. Proc Natl Acad Sci U S A 102:15545–50.

